# Synergistic Biophysics and Machine Learning Modeling to Rapidly Predict Cardiac Growth Probability

**DOI:** 10.1101/2024.07.17.603959

**Authors:** Clara E. Jones, Pim J.A. Oomen

**Affiliations:** Department of Biomedical Engineering, University of California, Irvine, CA 92697, USA; Edwards Lifesciences Foundation Cardiovascular, University of California, Irvine, CA 92697, USA

**Keywords:** Cardiac mechanics, Growth & remodeling, Biophysics modeling, Machine learning, Mitral regurgitation

## Abstract

Computational models that can predict growth and remodeling of the heart could have important clinical applications. However, the time it takes to calibrate and run current models while considering data uncertainty and variability makes them impractical for routine clinical use. This study aims to address this need by creating a computational framework to efficiently predict cardiac growth probability. We utilized a biophysics model to rapidly simulate cardiac growth following mitral valve regurgitation (MVR). Here we developed a two-tiered Bayesian History Matching approach augmented with Gaussian process emulators for efficient calibration of model parameters to align with growth outcomes within a 95*%* confidence interval. We first generated a synthetic data set to assess the accuracy of our framework, and the effect of changes in data uncertainty on growth predictions. We then calibrated our model to match baseline and chronic canine MVR data and used an independent data set to successfully validate the ability of our calibrated model to accurately predict cardiac growth probability. The combined biophysics and machine learning modeling framework we proposed in this study can be easily translated to predict patient-specific cardiac growth.

## 1 Introduction

Computational models have significantly advanced our understanding and management of heart diseases. These models, developed across various scales ^1^, encompass a broad spectrum of cardiac research areas, including hemodynamics ^2–4^, mechanics, ^5,6^, cell-signaling ^7^, and electrophysiology ^8,9^. With the increasing amount and quality of diagnostic data, as well as available computational resources, cardiac models now have the potential to be aimed towards providing a precision medicine approach ^1,10^. This innovation is poised to bring about a transformative shift in clinical practice, promising personalized treatment plans, enhanced diagnostic accuracy, and a deeper, more nuanced understanding of cardiovascular health.

While most current cardiac models can predict short-term outcomes such as QRS duration and ejection fraction, many are not necessarily predictive of long-term outcomes. The progression of heart disease is largely driven by growth and remodeling and therefore modeling these changes over time is imperative to fully understand and predict long-term cardiac disease outcomes. Several finite-element models have utilized changes in mechanics to predict observed trends in cardiac growth in response to classical cases of pressure ^11–14^ and volume overload ^12,13,15^. However, these finite element models are computationally expensive, especially when simulating months of cardiac growth and during model calibration, where numerous model evaluations are required to fit model parameters to match model outputs to experimental data. While computationally efficient cardiac growth models have recently been developed ^16,17^, the need for swift and precise models of long-term growth outcomes remains critical.

A further challenge in cardiac modeling is taking into account data uncertainty and variability. In the context of cardiac modeling, data uncertainty can arise from the imprecision in measuring patient-specific heart parameters such as end-diastolic volume, while variability is manifested through the natural differences in heart size, shape, or function across different subjects. Traditional calibration methods typically yield a singular set of fitted input parameters, rendering models calibrated in this manner incapable of accounting for data uncertainty and variability. This is particularly relevant for predicting cardiac growth, where uncertainty and variability may increase with time. Statistical methods such as Markov chain Monte Carlo have been used to address this issue ^18,19^, yet come with a high computational cost, making them cumbersome for cardiac growth models.

An alternative to traditional iterative model calibration methods is Bayesian history matching (BHM). Originating as an oil reservoir model calibration method in 1974^20^, BHM has since been used to calibrate computationally expensive models of galaxy formation ^21^, HIV epidemic progression ^22^, climate change in the Amazon rainforest ^23^, and, more recently, models of atrial electrophysiology ^24^ and ventricular mechanics ^25,26^. It uses a probabilistic approach to identify parameter combinations most likely to generate outputs consistent with the confidence interval (CI) of the observed data. It is typically augmented with a computationally efficient machine learning based emulator, or surrogate model, such as Gaussian process emulators (GPEs) ^27^, to accelerate model evaluations during the calibration process. GPEs have two advantages over other machine learning-based surrogate models such as neural networks that make them uniquely suited to cardiac modeling: they require fewer data to calibrate and return not only the predicted outcome but also its uncertainty inherent to the emulator.

The goal of this work is to create a computational framework to efficiently predict not just cardiac growth, but cardiac growth *probability*. We used our recently developed biophysics model ^17^ to rapidly simulate cardiac growth following mitral valve regurgitation (MVR), and BHM with GPEs for efficient calibration of the model parameters to align with growth outcomes within a 95*%* CI. Our approach involves two calibration steps to identify model parameters, similar to previous growth calibration work ^16,17^: first, adjusting cardiac and hemodynamic parameters for pre- and immediate post-MVR conditions, and then, calibrating growth parameters to match long-term changes. This dual-step calibration is novel in BHM model applications and is achieved by first optimizing cardiac and hemodynamic parameters before randomly sampling them to calibrate growth parameters. This method guarantees that our calibrated model precisely captures the progression of MVR from its onset through to chronic stages across varying degrees of initial disease severity. We first generated a synthetic data set with varying uncertainty (10 and 20% standard deviation) to assess the accuracy of our framework, and the effect of changes in data uncertainty on growth predictions. We then calibrated our model to match baseline and chronic canine MVR data, and used an independent data set to validate the ability of our calibrated model to accurately predict cardiac growth probability. A graphical presentation of the outline and calibration framework of this study is presented in Fig. 1.

**Figure 1:**
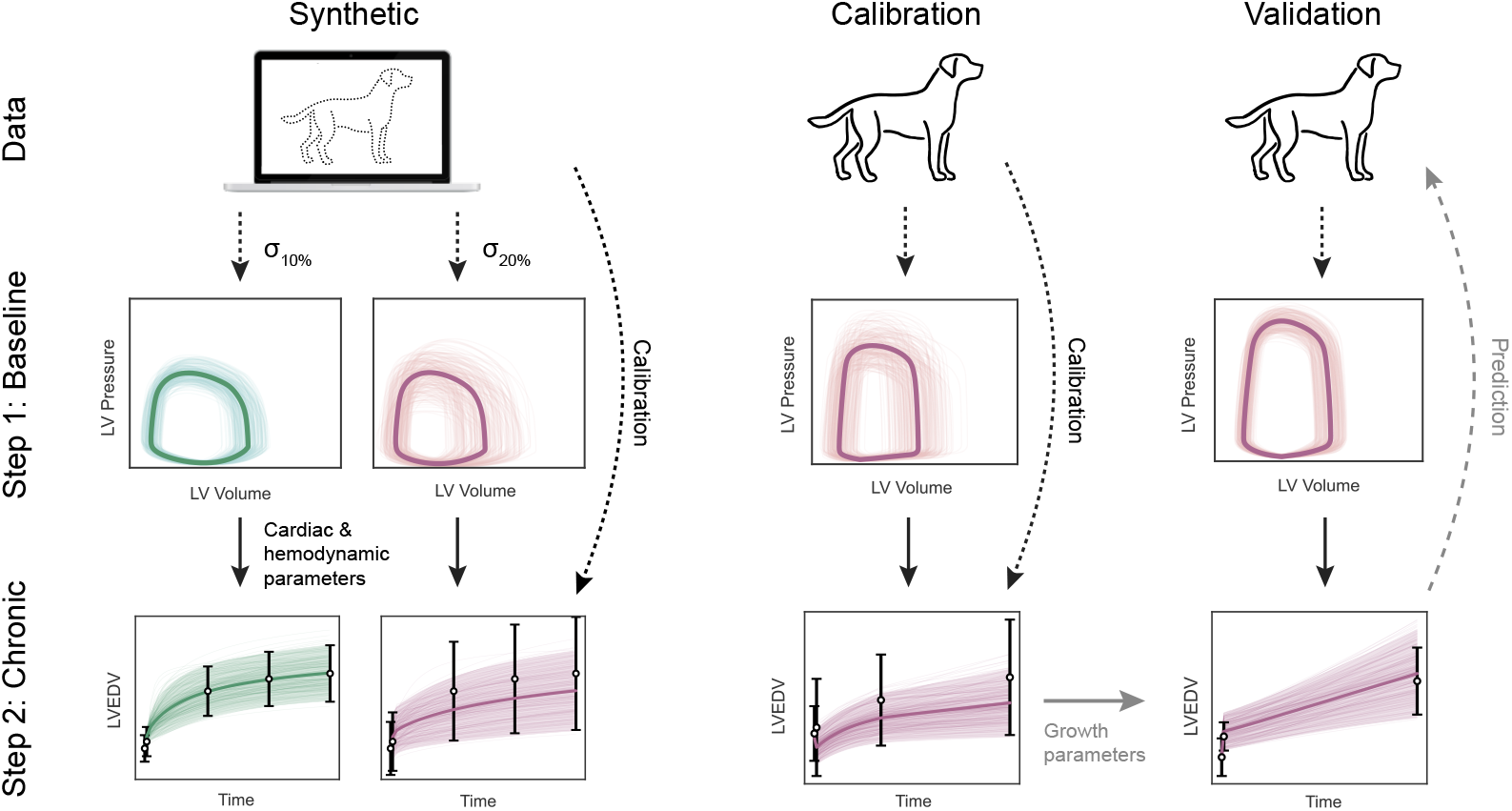
Graphical representation of the paper outline and calibration framework. The first step consists of using baseline pre- and post-MVR data (dotted lines) to calibrate cardiac and hemodynamic parameters. In the second step, the fitted parameter sets (solid lines) are randomly sampled during the calibration of the growth parameters to chronic data. We first assessed the accuracy of our method using synthetic data with varying uncertainty, then calibrated and validated the model using two independent canine MVR data sets.

## 2 Methods

### 2.1 Biophysics model

#### 2.1.1 Cardiac biomechanics

Our previously published model was utilized to simulate cardiac function and biomechanics ^17^. This approach uses the TriSeg method ^28^ to relate changes in ventricular blood volume to pressure. In brief, the left and right ventricles were modeled as three thick-walled semi-spherical walls: left (Lfw) and right (Rfw) free walls, and a shared septal wall (Sw). The spherical walls encapsulated the two ventricular cavities and coincided at a junction circle. The model schematic is shown in Fig. 2.

**Figure 2:**
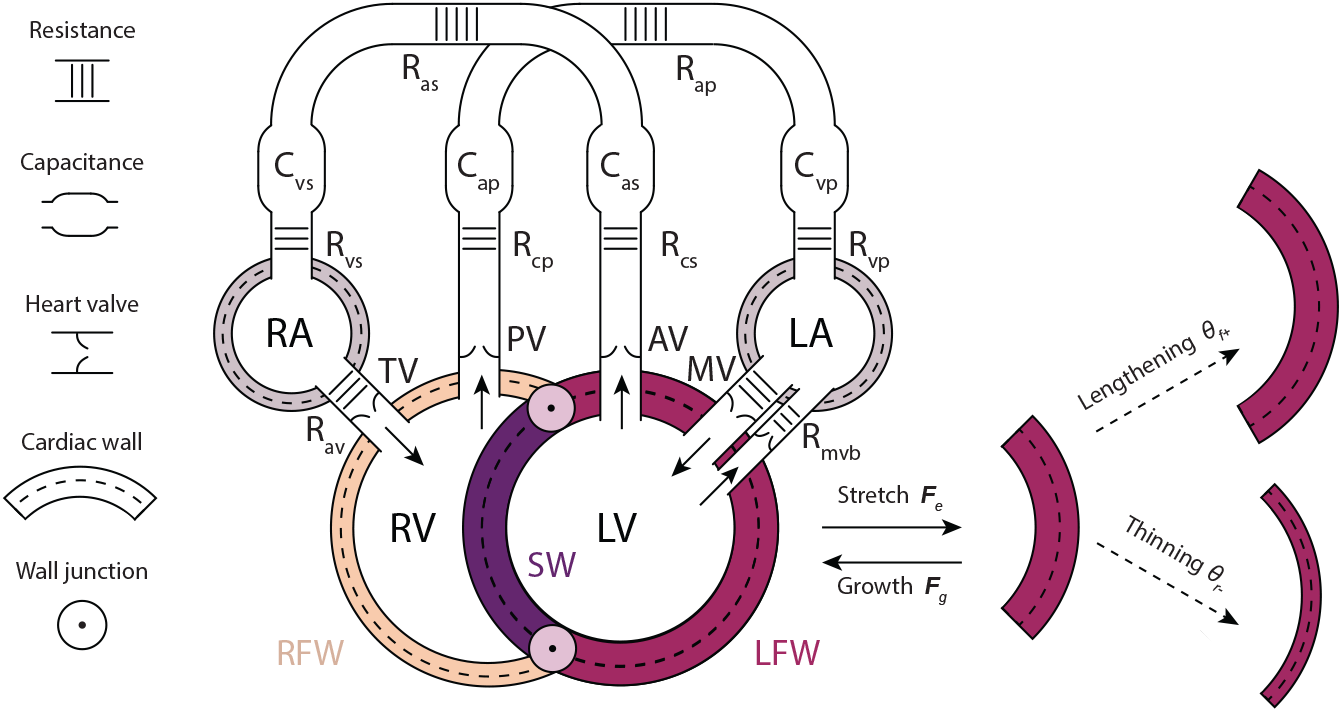
Schematic representation of the biophysics model. Cardiac mechanics were modeled using semi-spherical walls. Growth of the septal wall (SW) and left free wall (LFW) was determined using a transversely isotropic kinematic growth model driven by changes in circumferential and radial elastic stretch. The heart model was coupled to a lumped-parameter model to represent hemodynamics, where systemic and pulmonary arterial and venous flow was regulated using an electrical analog model of resistances, capacitances, and diodes. MVR was simulated by decreasing the mitral valve backflow resistance *R*_*mvb*_.

**Figure 3:**
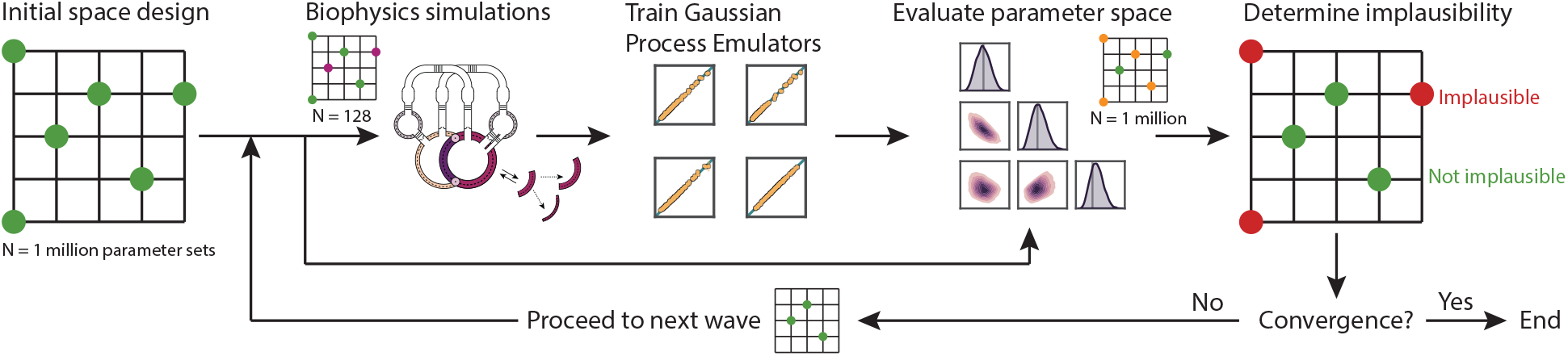
Schematic representation of the BHM algorithm. Starting with a parameter space of over one million points, a subset of 128 sets is utilized to run the biophysics model and subsequently train and validate GPEs on each model output. The full parameter space is then rapidly evaluated using the trained GPEs and parameters that lead to implausible solutions are discarded. The non-implausible parameters are passed on to the next wave, where this process repeats. By reducing the implausibility score with each subsequent wave, the parameter space progressively shrinks and the emulators get trained better in the emerging region of interest.

The ventricular geometry was assumed to be axisymmetric around an axis perpendicular to this junction circle. Mechanics were calculated at the midwall surface area *A*_m_, defined as the surface that divides the wall volume *V*_w_ in two equal volumes. We assumed a transversely isotropic fiber distribution oriented along *A*_*m*_. Sarcomere elastic fiber stretch *λ*_f_ was calculated as the relative change in midwall surface area *A*_m_ compared to the midwall surface area in the unloaded reference configuration *A*_m,ref_, and radial stretch by assuming incompressibility:

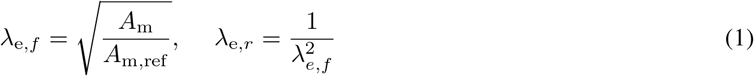

Fiber stress was a function of fiber stretch *λ*_e,f_ and activation timing and decomposed into a passive (*σ*_f,pas_) and active (*σ*_f,act_) component:

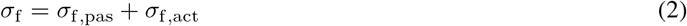

Passive stress was calculated using an exponential co stitutive m del from ^17^:

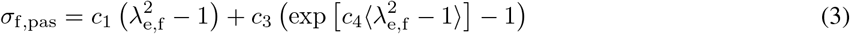

with *c*_1_, *c*_3_, and *c*_4_ material properties, and the Mackaulay brackets ⟨°⟩ enforcing fiber nonlinear stress only when at tension. Active stress was calculated using the Hill-type single fiber active contraction model from ^4^. In brief,

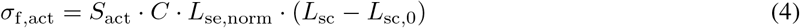

where *S*_*f*,act_ is a linear coefficient scaling how much active stress the sarcomere can generate; *C* a state variable of contractility as a function of time and fiber stretch *λ*_*e,f*_; *L*_se,norm_ the length if the normalized series elastic element in the Hill-type model; *L*_sc_ and *L*_sc,0_ the sarcomere contractile element length at the current and unloaded state, respectively. The midwall tension *T*_*m*_ in each semi-spherical wall was then approximated using ^29^:

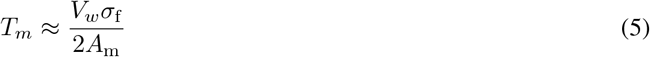

and an iterative local Newton scheme was employed to achieve a balance of tension between the three walls at the junction points. Convergence was established by adjusting the height of the junction circle and deflection of the Sw relative to the left and right free walls. Finally, the pressure *p* in each cavity was computed using Laplace’s law and the midwall radius *r*_*m*_:

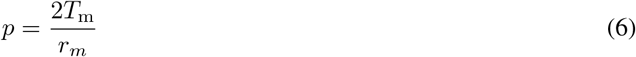

#### 2.1.2 Hemodynamics

To model hemodynamics, we employed a lumped-parameter model of the circulation based on Santamore & Burckhoff ^3^. Changes in blood volume and pressure in the arterial and pulmonary arteries and veins, as well as left and right ventricles and atria were modeled using an electrical circuit analogy. The total blood volume contained in the circulation was functionally divided into two parts: unstressed and stressed blood volume (SBV). Unstressed blood volume was defined as the maximum blood volume that can fit within a compartment without causing the pressure to rise form 0 mmHg, here assumed to remain constant. The pressure in each vascular compartment was related to the compartment’s (stressed) volume *V* and capacitance *C*,

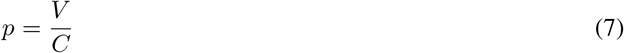

where the flow *q* between two adjacent compartments was determined by the difference in pressure

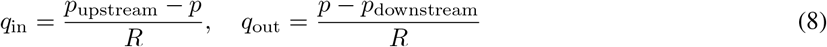

where *R* is the resistance between the two compartments. The total flow was then determined by the difference in pressure between adjacent vascular compartments.

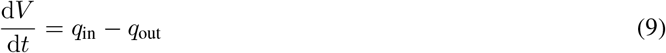

Pressure-sensitive diodes represented heart valves, where flow to the downstream compartment was only allowed when the pressure in the upstream compartment exceeded the pressure in the downstream compartment. Thus, we have a system of 8 (4 vascular and 4 cardiac) differential equations for *V* (*t*), which we solved over the cardiac cycle using the fourth-order Runge-Kutta method. MVR was implemented by allowing flow from the LV back to the left atrium, proportional to a backflow resistance, *R*_*mvb*_:

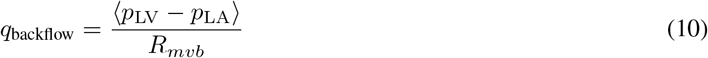

with the Mackaulay brackets ⟨□⟩ ensuring regurgitation only occurring when the pressure in the LV exceeded pressure in the left atrium. In healthy hearts, we set *R*_*mvb*_ → ∞ to prevent regurgitation.

#### 2.1.3 Growth model

To simulate left ventricular growth, we employed a mechanics-driven cardiac growth model based on the kinematic growth framework ^30^. The total deformation gradient tensor ***F*** was multiplicatively decomposed into an elastic component ***F***_*e*_ and a growth component ***F***_*g*_:

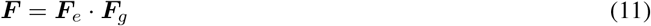

The elastic tensor is composed of the fiber and radial stretch from the mechanical model of the heart (Section 2.1). To model dilation and thickening, and considering that in our model the sarcomere fibers are distributed isotropically along the myocardial surface, we chose a transversely isotropic growth tensor:

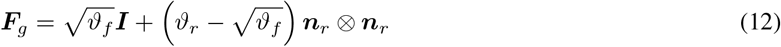

where *ϑ*_*f*_ and *ϑ*_*r*_ are so-called growth multipliers in the fiber and radial directions, *n*_*r*_ is the unit vector in the radial direction, and ***I*** is the identity tensor. Note that when all growth multipliers equal 1 then ***F***_*g*_ = ***I***. This formulation allows for independent growth in the fiber and radial directions. The evolution of the growth multiplier in both the fiber and cross-fiber directions was governed by a limiting function *k* and a stimulus function *ϕ*

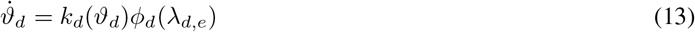

with *d* = *f, r*, and its evolution between time points *t*^*n*−1^ and *t*^*n*^ given by the finite difference

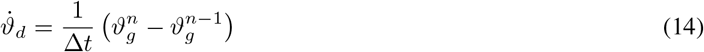

The weighing function *k* limits growth towards a certain maximum amount of growth 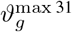:

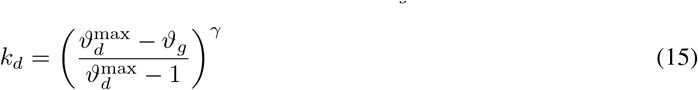

where the time constant *τ* represents the sarcomere deposition time and *γ* represents the sarcomere deposition non-linearity, here set to 2, and *ϑ*^max^ represents the cardiomyocyte growth limit. We set this to 1.5 for growth and 0.75 for reverse growth. Although there is no known study on the maximum and minimum cardiomyocyte dimensions, we assume there is a physiological limit to cardiomyocyte growth based on literature studying cardiomyopathic cardiomyocyte dimensions, with the maximum and minimum growth limits we set encompassing the data from this study ^32^. The stimulus functions were adapted from Göktepe et al. ^33^, where dilation in the fiber direction *ϕ*_*f*_ growth was driven by the difference between the current stretch in the fiber direction *λ*_e,f_ (Eq. 1) and a homeostatic stretch setpoint 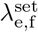:

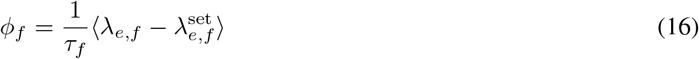

with *τ*_*f*_ a time constant for the fiber sarcomere deposition rate. Radial thinning is driven by the difference between the current stress in the fiber direction *σ*_*f*_ (Eq. 2) and a homeostatic stress setpoint *σ*^set^:

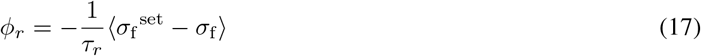

with *τ*_*r*_ a time constant for the radial sarcomere degradation rate. Note that the Mackaulay brackets ⟨°⟩ were used the to enforce fiber lengthening and radial thinning only, as these are the only growth patterns observed in volume overload so it would not be possible to calibrate our model parameters to fiber shortening and radial thickening. Volume and wall area are then updated at each time point using the growth multipliers. Since our fiber distribution is isotropic, the growth multiplier *ϑ*_*f*_ represents area growth and therefore the change in wall area, 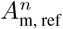 at time point *t*^*n*^ can be found through:

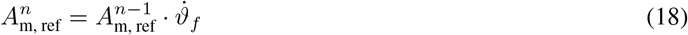

Similarly, the wall volume, 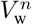 is found via the total change in volume during the current growth increment:

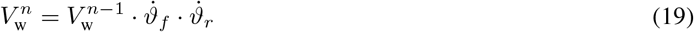

### 2.1 Model calibration

#### 2.2.1 Gaussian Process Emulation

We used Gaussian Process Emulators (GPEs) as a surrogate for our biophysics models, which allows for accelerated evaluation of a large parameter space. This was done using the GPyTorch package in Python ^34^. Before emulator training, all data is normalized. A GPE *f* was generated to estimate each model output,

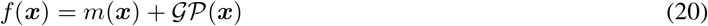

with *x* a vector of model inputs (see Section 2.3), *m* a linear mean function, and GP a Gaussian process with a radial basis function (RBF) as covariance matrix and Gaussian noise with variance 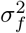. The hyperparameters of each GPE were optimized using the Adam optimizer with a learning rate of 0.1 and 100 iterations to maximize the marginal log likelihood, defined in GPyTorch ^34^ as

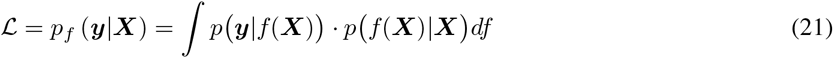

where ***y*** is a vector with simulated values and ***X*** a matrix containing model input vectors *x*. The first term *p*(*y*|*f* (***X***)) represents how likely it is that the simulated outputs match the emulated outputs, while the second term *p*(*f* (***X***)|***X***) represents the certainty of the emulator. We used a random selection of 80% of the filtered simulated results to train the model and 20% for validation. The *R*^2^ score between the simulated and emulated results was calculated. On the total set of simulations run to train and test the emulators, a sequence of mean absolute deviation and Mahalanobis distance multivariate filtering was used to exclude outliers from the simulated outputs allowing for more robust GPE training. Additionally, parameter sets yielding non-physiological results (e.g. mean arterial pressure *>* 250 mmHg) were excluded.

#### 2.2.2 Bayesian History Matching

We calibrated our biophysics model to match synthetic and experimental data (See Section 2.3) using Bayesian History Matching (BHM) augmented with GPE surrogate models. This is a statistical method that iteratively refocuses the model’s parameter space in ‘waves’. The first wave starts with a large parameter space (*n* = 2^20^ = 1, 048, 576) generated from a Sobol sequence as a space-filling design to cover the entire parameter space as efficiently as possible.

An additional sampling sequence was performed on this space using the *diversipy* library to select *n* = 2^7^ = 128 points to perform biophysics simulations. The results from these simulations were used to train and validate the GPEs as described in Section 2.2.1. The trained GPEs were utilized to evaluate the entire parameter space. We used the implausibility measure ^21,35^ to evaluate whether each parameter set ***x*** resulted in an acceptable match between the emulator output and the data, taking into account emulator and data uncertainties. For a single output *i* the implausibility was calculated as

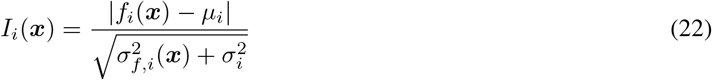

where *f*_*i*_ is the emulated value of output *i* (Eq. 20), *µ*_*i*_ the mean target data, and 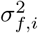 and 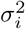 the variance of the emulator and data, respectively. To assess the implausibility of all outputs, we use the maximum implausibility *I*(*x*) = argmax_*i*_(*I*_*i*_(***x***)), constituting a worst-case scenario. Parameter sets that satisfy *I < I*_threshold_ are considered non-implausible and form the parameter space for the next BHM wave. This process was repeated over several waves, with the threshold lowered incrementally until the parameter space has converged. As the parameter space shrinks throughout subsequent waves, the biophysics simulation data becomes denser in that area of focus, thus reducing emulator variance and causing more points to be deemed implausible. In this study, we chose 2 as our final implausibility score to ensure all points were within two standard deviations, i.e. a 95% CI of the data. 256 parameter sets from the final parameter space were randomly chosen to run posterior biophysics simulations. If fewer than 50,000 points remained in the parameter space at the end of each wave, the cloud method ^24^ was used to generate additional data points.

#### 2.2.3 Two-step approach

Mechanics-driven growth simulations typically require three simulation stages: healthy pre-disease, acute post-disease, and chronic post-disease cardiac growth. The extent of growth depends on a combination of the severity of the disease (here: the degree of MVR), i.e. the difference at baseline between the healthy and diseased states, and the potential of cardiomyocytes to grow in response to the change in mechanics imposed by the disease. Therefore, parameters required to simulate long-term cardiac growth need to be calibrated independently from the cardiac and hemodynamic parameters that are required to simulate the two baseline stages. To achieve this, we employed two BHM steps to identify model parameters: (1) calibrating cardiac and hemodynamic parameters to match the baseline pre- and immediate post-MVR conditions and (2) calibrating growth parameters to match long-term changes (Fig. 1). In the second step, we randomly sampled cardiac and hemodynamic parameters from the first step to propagate parameter uncertainty and variability throughout the growth simulation.

#### 2.2.4 Model inputs and outputs

The baseline and growth parameter values and their bounds for calibration can be found in Table 1 and were based on previous work from Witzenburg & Holmes ^16^ and Oomen et al. ^17^. The cardiac and hemodynamic parameters used to calibrate model baseline pre- and post-MVR in BHM step 1 were

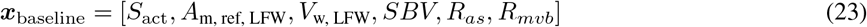

and the growth parameters used to calibrate chronic MVR in BHM step 2 were

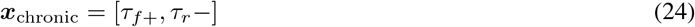

**Table 1:** Biophysics model input parameter used for calibration.

| Stage | Parameter | Description | Equation | Range | Synthetic value | Units |
| --- | --- | --- | --- | --- | --- | --- |
| Baseline | $S_{act}$ | Active stress coefficient | 4 | 0.050 – 0.200 | 0.150 | MPa |
| | $c_3$ | Passive stress coefficient | 3 | 5e-6 – 5e-3 | 5.23e-4 | - |
| | $A_{m, ref, LFW}$ | Left free midwall reference area | 1 | 4,000 – 10,000 | 6,000 | mm<sup>2</sup> |
| | $V_{w, LFW}$ | Left free wall volume | 5 | 40,000 – 70,000 | 40,800 | mm<sup>3</sup> |
| | $SBV$ | Total stressed blood volume | 7 | 300 – 1,500 | 1,000 | mL |
| | $R_{as}$ | Systemic arterial resistance | 8 | 0.3 – 5.0 | 1.1 | mmHg s mL<sup>-1</sup> |
| | $R_{m vb}$ | Mitral valve backflow resistance | 10 | 0.05 – 2.0 | 0.3 | mmHg s mL<sup>-1</sup> |
| Chronic | $\tau_{f+}$ | Fiber lengthening time constant | 16 | 0.1 – 50 | 1.0 | - |
| | $\tau_{r-}$ | Radial thinning time constant | 17 | 0.1 – 50 | 5.0 | - |

Based on the experimental data reported in the canine MVR studies we used for model calibration and validation (see Section 2.3), the following data means were used to calibrate the model baseline (Eq. 22):

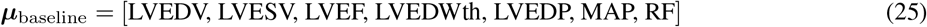

where LVEDV is LV end-diastolic volume, LVESV LV end-systolic volume, LVEF LV ejection fraction, LVEDWth end-diastolic left free wall thickness, LVEDP LV end-diastolic pressure, MAP mean (systemic) arterial pressure, and RF regurgitant fraction. Again, based on data availability, the chronic MVR response of the model was calibrated to match the following means:

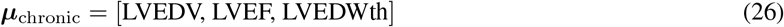

To assess each input parameter’s contribution to the model outputs’ variance, we performed a Sobol sensitivity analysis using the Saltelli method ^36^. This global first-order sensitivity analysis was implemented in the SALib Python package ^37^. Emulators trained during the first BHM wave were used to accelerate the otherwise computationally expensive analysis. Parameters that contributed less than 95% of the total variance were excluded from the fitting procedure.

### 2.3 Accuracy, calibration, and validation

We first assessed the accuracy of our two-step BHM method to calibrate a mechanics-based growth model by matching synthetic data that we generated using our biophysics model (Fig. 1). The data was generated by manually setting model parameters to result in simulations representative of MVR in canines ^38,39^. Model outputs were used as ‘mean’ outcomes (Eqs. 25 and 26). To assess the impact of data uncertainty and variability on calibration accuracy, two synthetic data sets were generated with standard deviations of 10% and 20%, i.e. *σ* = 0.1*µ* and 20% of the mean *σ* = 0.2*µ*, resp. We will denote these cases as *σ*_10%_ and *σ*_20%_. To test whether the optimized growth parameters were independent of the baseline parameters, we randomly redistributed the optimized baseline and growth parameters and re-evaluated the growth outcomes.

Next, we tested the real-world applicability of our method by calibrating our model to previously published data on chronic canine MVR by Kleaveland et al. ^38^ (Fig. 1). The same model parameters and outputs were used as in the synthetic cases. Finally, we validated the ability of our framework to predict growth probability independent of baseline calibration. This was achieved by fitting the cardiac and hemodynamic parameters of our model to match an independent chronic MVR study by Nakano et al. ^39^, followed by a prediction of cardiac growth by combining 256 remaining cardiac and hemodynamic parameters sets with a random sampling of 256 growth parameter sets from the calibration study.

## 2 Results

### 3.1 Demonstration of BHM utilizing synthetic dataset with varying uncertainty

We successfully calibrated our biophysics model to the synthetic baseline and chronic MVR data using our two-step BHM approach. The entire two-step calibration process took less than 10 minutes on a desktop PC with an Apple M1 Max processor, using 9 parallel processors. All model parameters converged to unimodal distributions centered around the ground truth mean. As expected, matching the *σ*_10%_ case resulted in parameter distributions with less variance compared to the *σ*_20%_ case. The growth time constants in particular had considerably larger variance towards the upper parameter limits (i.e. slower growth) for the *σ*_20%_ case (Fig. 4b,c). From the initial 2^20^ = 1, 048, 576 baseline parameter sets, only ∼7,300 (0.7*%*) remained for the *σ*_10%_ case, and ∼8,500 (0.8*%*) for the *σ*_20%_ case. More parameter sets remained from the initial growth parameter sets: 36,000 (3.6*%*) for the *σ*_10%_ case, and 73,000 (7.3*%*) for the *σ*_20%_ case. Emulators were highly accurate, with *R*^2^ scores of emulated vs. simulated results at least 0.96 for all emulators (Appendix Fig. A1).

**Figure 4:**
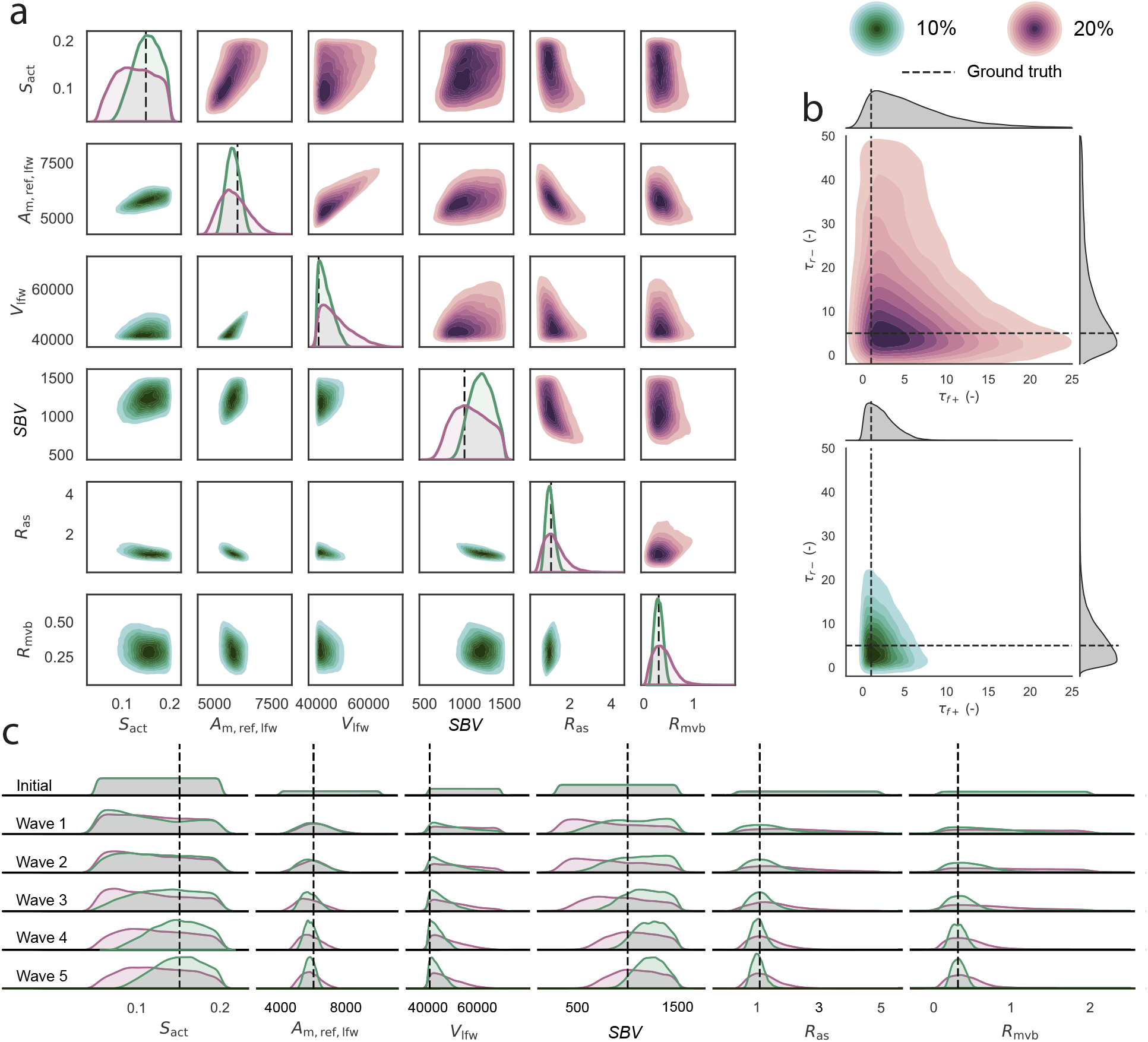
Optimized input parameter distributions. (a) Multidimensional cardiac and hemodynamic optimized parameter spaces found during BHM step 1 of the *σ*_10%_ (green shades, lower triangle) and *σ*_20%_ (purple shades, upper triangle) synthetic cases. (b) Optimized growth parameter spaces for the two synthetic cases. (c) Convergence of the cardiac and hemodynamic parameters throughout the 5 BHM waves. All model parameters converged to unimodal distributions centered around the ground truth mean (dashed lines), indicating that unique solutions were found for all parameters, but the *σ*_10%_ resulted in distributions with less variance than the *σ*_20%_ case.

Before starting the fitting procedure, the Sobol sensitivity analysis revealed that the passive constitutive parameter *c*_3_ was not contributing significantly to the variance of the model outputs (Fig. 5), so this parameter was omitted from the matching pipeline and set constant to match experimental data ^40^. Left free wall midwall reference area *A*_m,ref_ was the parameter that contributed most to the model output variance.

**Figure 5:**
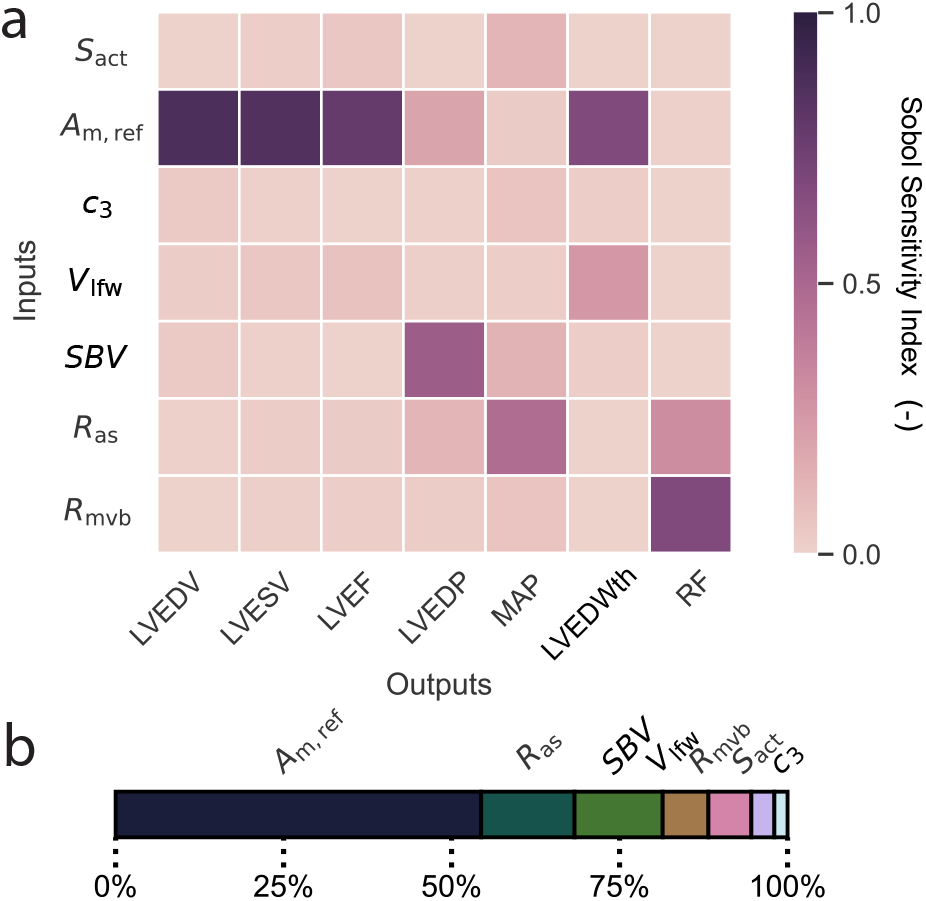
Global sensitivity analysis. (a) Total effects of cardiac and hemodynamic model input parameters (vertical axis) on the baseline outputs (horizontal axis), where a high first-order Sobol index indicates a larger sensitivity. (b) Normalized total sensitivity of each model parameter where each output’s sensitivity adds up to 1, with *A*_m,ref_ the most important parameter and *c*_3_ the least important one, which was thus excluded from the calibration process.

The calibrated baseline simulations produced outputs mostly within 95% CIs of the synthetic data (Fig. 6). All baseline (Fig. 6a) and acute (Fig. 6b) output distributions for the *σ*_10%_ case were unimodal and symmetric with the means centered around the ground truth mean for all outputs except LVEDWth, which was likely due to the chosen lower bound for the left free wall volume being close to the target value (Table 1). In contrast, the *σ*_20%_ case output distributions were skewed and not always aligned with the ground truth mean. For both cases, the resulting pressure-volume loops resulted in expected behavior for MVR, but with increased variance for the *σ*_20%_ case: widening of the loop, reduction in maximum pressure, and loss of isovolumetric contraction and relaxation (Fig. 7).

**Figure 6:**
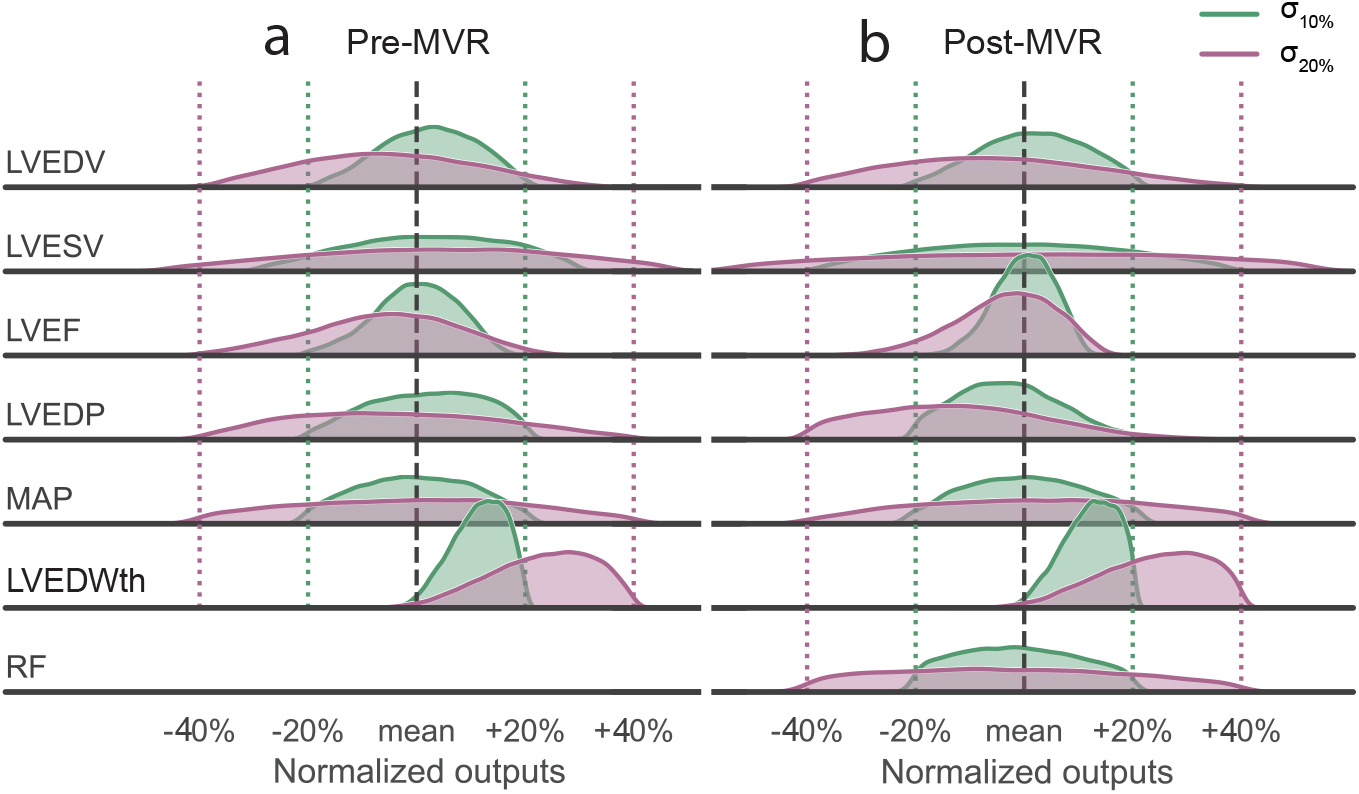
Comparison of calibrated baseline model outputs to synthetic data. Calibrated baseline pre-(a) and post-MVR (b) model outputs were normalized to the data means (dashed lines). Simulated outputs of both the *σ*_10%_ (green) and *σ*_20%_ (purple) cases were within the data 95% CIs (dotted lines), which equal 20% and 40% of the mean for the *σ*_10%_ and *σ*_20%_ cases, resp. All output distributions unimodally distributed and, except for LVEDWth, centered around the mean. RF was not fitted for the pre-MVR case since we assumed no MVR at that stage and LVEDWth was not reported in the validation study.

**Figure 7:**
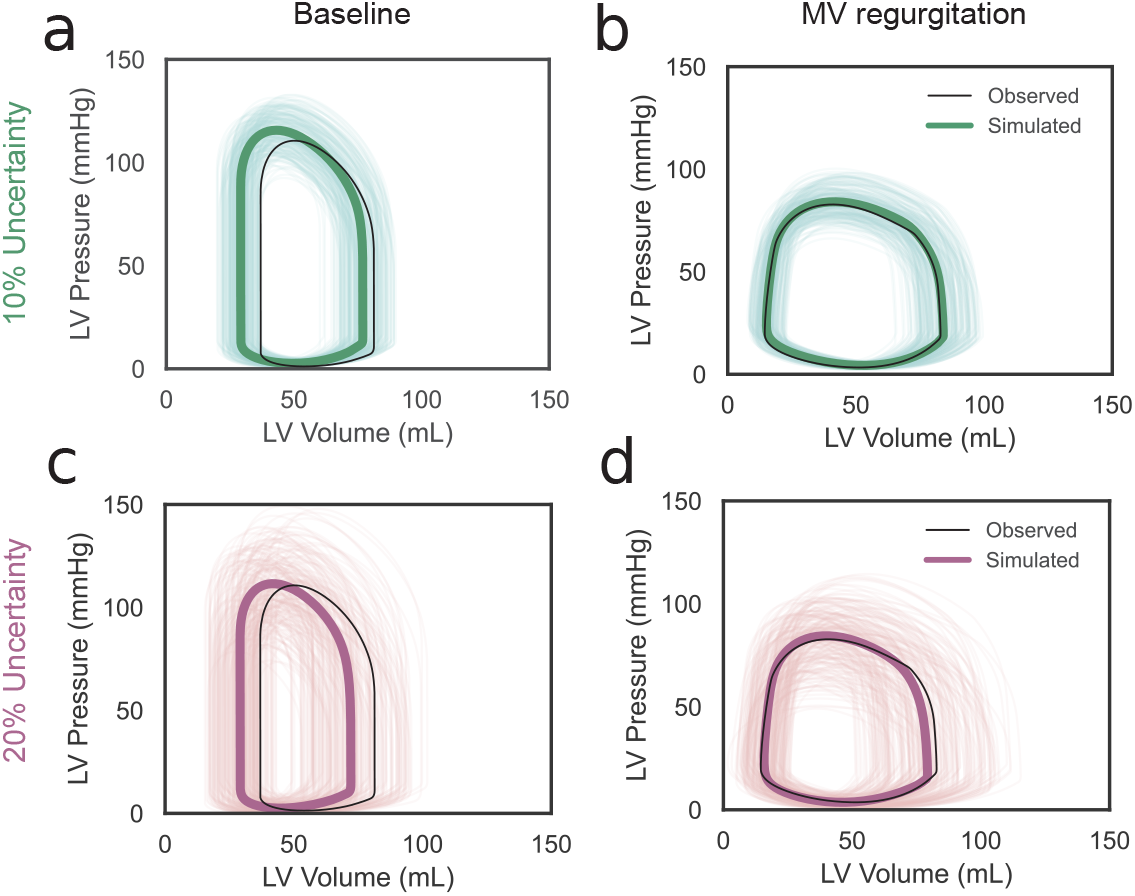
Comparison of LV pressure-volume loops to synthetic data. Pressure-volume loops of the *σ*_10%_ (a,b, green) and *σ*_20%_ (c,d, purple) calibration cases closely matched the synthetic data (black lines). The individual calibrated simulation results are represented by thin lines, their average values by thick lines, and the 95% CI by shaded regions. The *σ*_20%_ case resulted in higher pressure-volume variance than the *σ*_10%_ case and a overestimation of the pre-MVR mean volumes.

The growth simulations from the calibrated biophysics model also produced outputs within 95% CIs of the synthetic data (Fig. 8). The biophysics model correctly matched the increase in LVEDV and decrease in LVEF and LV wall thickness over the course of 90 days. The matched means and variance of the *σ*_10%_ case aligned well with the ground truth. However, the simulated LVEDV means for the *σ*_20%_ was lower than the ground truth mean, with variance only increasing towards the lower range of the CI. Conversely, LVEF was overestimated, with the variance increasing towards the upper range of the CI. Simulated changes in wall thickness were similar in the *σ*_10%_ and *σ*_20%_ cases. Randomly recombining the baseline and growth parameters resulted in identical outcome distributions, indicating that the growth parameters were fitted independently of the baseline parameters (Appendix Fig. B1).

**Figure 8:**
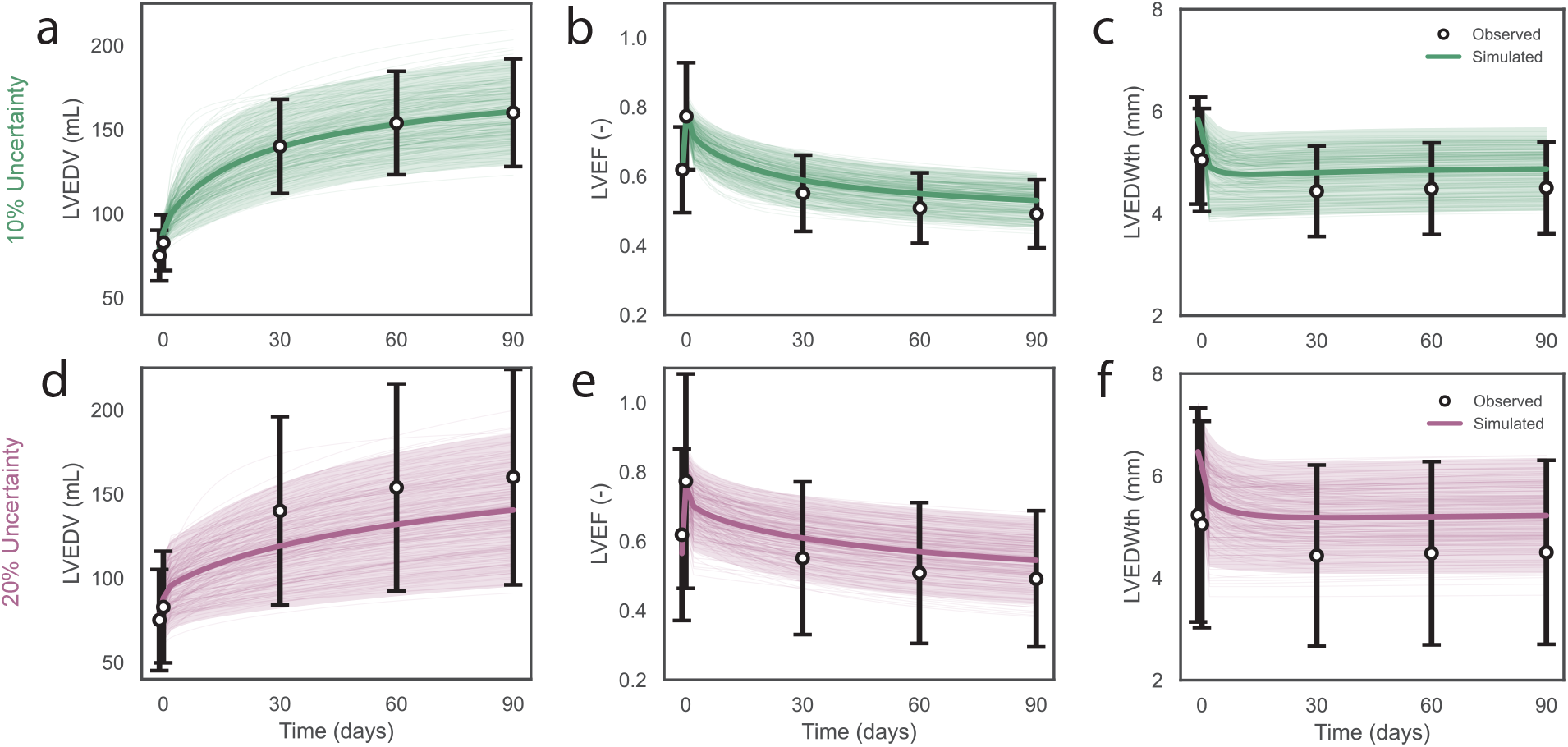
Comparison of growth simulations to synthetic data. Simulated chronic post-MVR LVEDV (a,d), LVEF (b,e), and LVEDWth (c,f) were within the 95% CI of the synthetic data (black whiskers) for both the *σ*_10%_ (a-c, green) and *σ*_20%_ (d-f, purple) cases. The individual calibrated simulation results are represented by thin lines, their average values by thick lines, and the 95% CI by shaded regions. The means of the *σ*_10%_ case closely matched the data, whereas LVEDV was overestimated and LVEF and LVEDWth were underestimated for the *σ*_20%_ case together with increased variance.

### 3.2 Growth model calibration and validation with experimental data

Similarly to the synthetic case, we successfully calibrated our biophysics model to match the real-world baseline data from the calibration ^38^ and validation ^39^ studies (Fig. 9). The posterior distributions were unimodal and mostly within 2 standard deviations of the experimental data. All distributions were centered around the mean except for LVEDV and LVEDWth of the calibration baseline post-MVR case.

**Figure 9:**
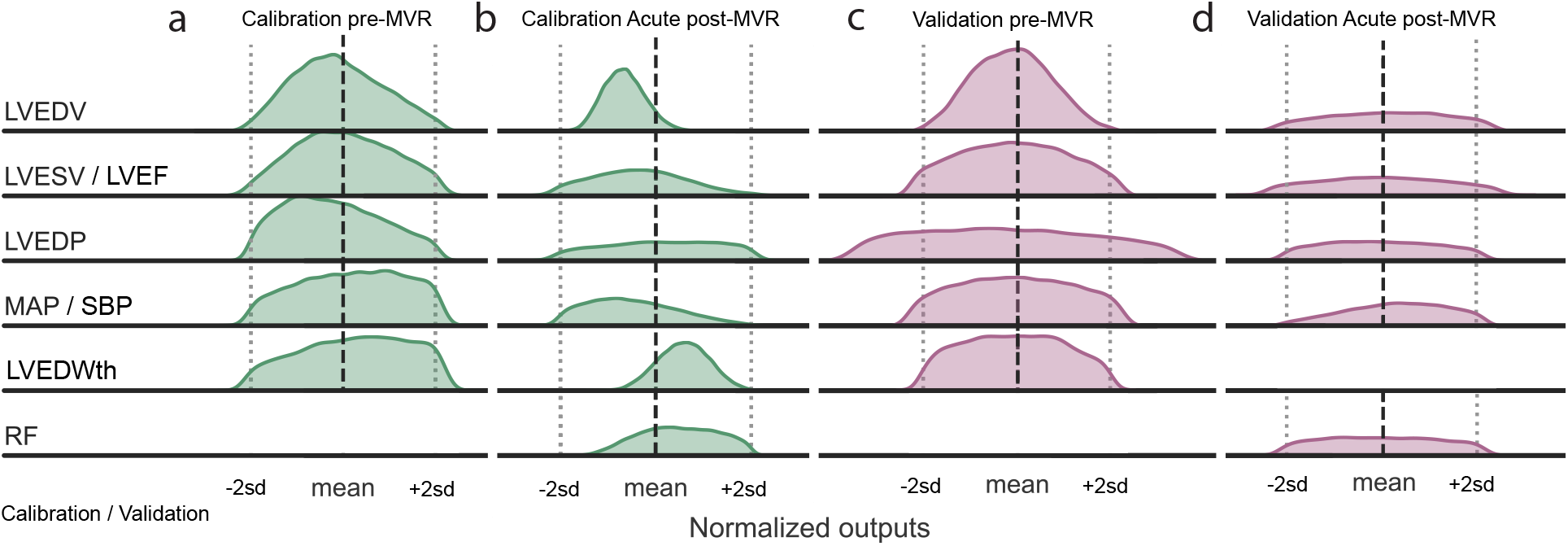
Comparison of calibrated baseline model outputs to canine data. Calibrated baseline pre-(a, c) and post-MVR (b, d) model outputs were normalized to the data means (dashed lines). Simulated outputs of both the calibration (a,b, green) and validation (c,d, purple) cases were within the data 95% CIs (dotted lines), except for LVEDP of the validation case. All outputs distributions unimodally distributed and, except for LVEDV and LVEDWth for the calibration case, centered around the mean. RF was not fitted for the pre-MVR case since we assumed no MVR at that stage.

We successfully optimized the growth parameters to simulate chronic changes in LV dimensions. The simulated growth was mostly within two standard deviations of the observed calibration data (Fig. 10), albeit with slightly underestimated means. The optimized growth parameters from the calibration case were then randomly combined with the optimized cardiac and hemodynamic parameters of the validation case to predict growth for the validation case ^39^. The predicted mean change and distribution of LVEDV aligned well with the experimental data, with most posterior simulations falling within the reported 95% CI. LVEF was slightly overestimated, with the mean and most model predictions within the reported CI. However, wall thinning predictions were underestimated, with the mean and most model predictions falling below the reported data interval.

**Figure 10:**
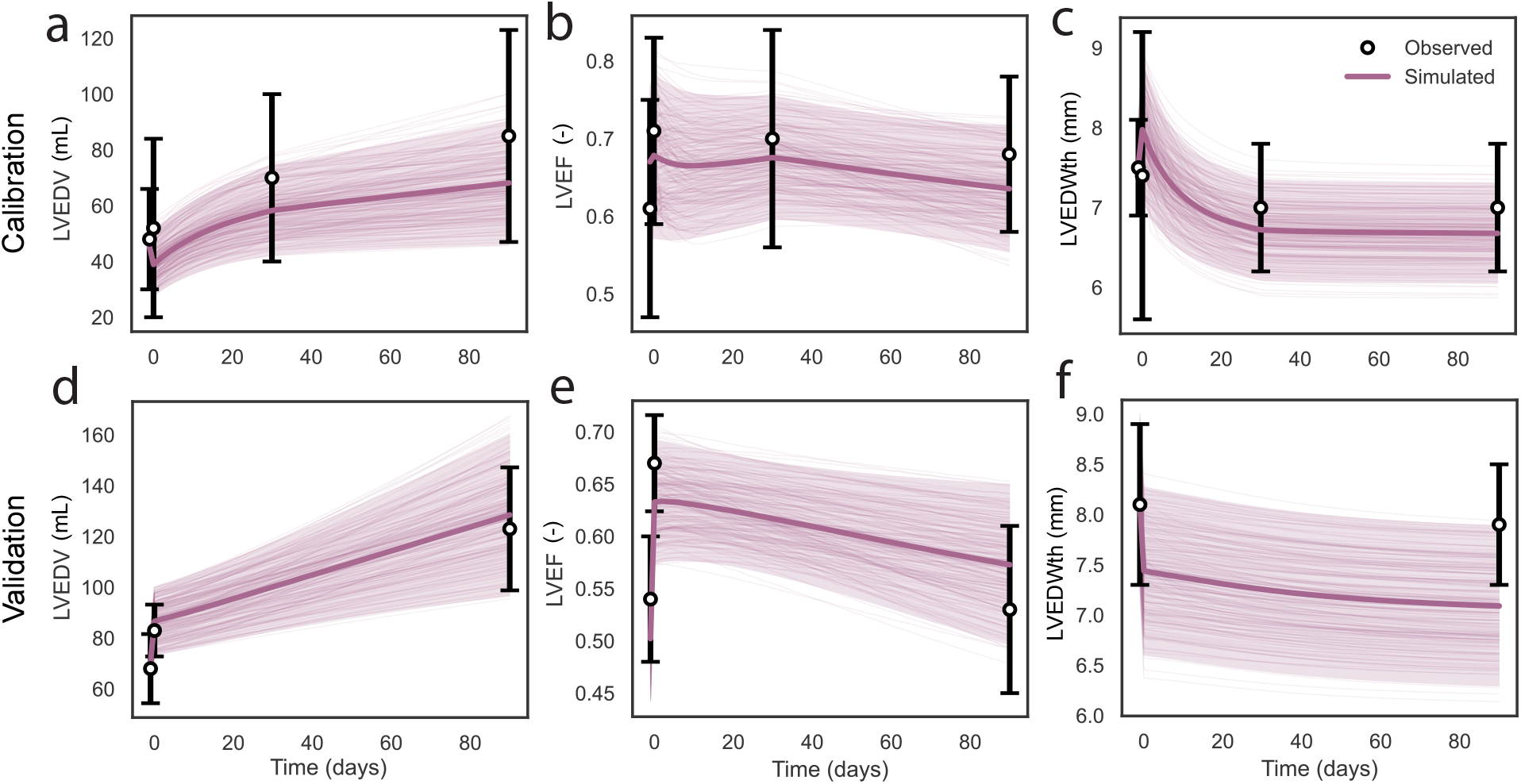
Canine growth calibration and predictions. Calibrated chronic post-MVR LVEDV (a), LVEF (b), and LVEDWth (c) matched 95% CI of the canine data. The individual simulation results are represented by thin lines, their average values by thick lines, and the 95% CI by shaded regions. The calibrated growth parameters were then used to predict growth for the independent validation case. The predicted mean and 95% CI of LVEDV (d) closely matched the canine data, while predicted LVEF (e) was slightly overestimated and LVEDWth (f) underestimated.

## 4 Discussion

This study aimed to create a computational framework to efficiently predict cardiac growth probability. Using techniques previously used in other fields such as BHM and GPE, we created a machine learning-based two-step method to rapidly calibrate a cardiac biophysics model to simulate cardiac growth probability during MVR. We first tested the feasibility and accuracy of our method using synthetic data generated by our biophysics model, including the effect of data uncertainty on growth outcome (Figs. 4-8). We then predicted 3 months of growth in canines ^39^ using growth parameters calibrated to an independent data set ^38^ (Figs. 9, 10). The results confirm that our model can be rapidly calibrated and then used to predict growth in a different case, laying the groundwork for patient-specific model calibration.

### 4.1 Comparison with prior work

The main advantage of BHM over iterative fitting methods such as least squares and gradient descent is that it accommodates biophysics models to predict a range of probable outcomes instead of a single outcome. This is because BHM employs an implausibility criterion using Bayes’ theorem (Eq. 22) to identify all model parameters that result in model results within a certain statistical interval, in our case, a 95% CI. In contrast, iterative methods identify a single minimal value of an objective function to minimize the difference between the model-estimated value and the data. Thus, BHM can identify growth probability, instead of yielding only a single simulation result that is the closest to the data mean. This advantage of BHM over iterative fitting can be appreciated when comparing our results to the recent study by Witzenburg & Holmes ^16^. They were able to, for the first time, predict LV growth and hemodynamic following pressure overload, volume overload, and myocardial ischemia using a single parameter set identified used the least squares method. While their model predictions were within the experimental standard deviation, they did not identify the full range of potential growth outcomes. Here, we used the same canine volume overload data sets as in Witzenburg & Holmes to calibrate and validate our model and demonstrated that we can indeed predict the full range of possible growth outcomes (Fig. 10). Note that we used a different growth model formulation than Witzenburg & Holmes (Section 2.1.3): the change of the cardiac growth tensor in their formulation was multiplicative based on the work from Kerckhoffs et al. ^15^, whereas ours, based on prior work from Göktepe et al. ^12^, evolves additively, which results in more stable outcomes. This increased stability was important in the context of this work since, especially during the first BHM wave, the wide range of growth parameters that we evaluated resulted in unstable growth outcomes when using the multiplicative Kerckhoffs formulation.

The first simulations of cardiac growth that propagated data uncertainty were recently performed by Peirlinck et al. ^19^. They used Bayesian inference and Gaussian process regression to quantify the effect of changes in stretch on myocyte growth following MVR in porcine. They used a similar mechanics-based growth relationship as in our model (Eqs. 16, 17) to drive myocyte growth, but did not formally calibrate any of the growth parameters as we did in this study using BHM.

While BHM has been widely applied in other engineering fields, it has only recently been adopted for cardiovascular engineering. Coveney & Clayton optimized an atrial electrophysiology model to simulate action potentials of atrial fibrillation patients ^24^. Longobardi et al. optimized a biventricular electromechanical model to match in vivo rat data before and after left ventricular pressure overload ^25^. The calibration method described in this paper is based on these two studies, with a few notable differences. First, we used a two-step approach, where we first identified pre- and post-MVR hemodynamic and cardiac parameters and then sampled from these parameters to identify growth parameters (Section 2.2.3). This was crucial to identify growth parameters that could predict changes in LV dimensions within the full range of change between pre- and post-MVR baseline parameters (Appendix Fig. B2). Second, we chose a lower implausibility criterion than in the previous studies. Both Coveney & Clayton and Longobardi et al. used a final implausibility criterion of *I <* 3 using the Pukelsheim 3-sigma rule, which states that 95% of all observed values are within three standard deviations when assuming a symmetric unimodal distribution ^41^. However, we chose to use *I <* 2 to maintain all model predictions within two standard deviations according to the empirical 68-95-99 rule.

### 4.2 Synergy of biophysics and machine learning modeling

As discussed above, the main strength of BHM is that it enables the identification of parameters that can predict the full range of possible growth outcomes conditioned on available hemodynamic data. This does, however, require substantially more model evaluation evaluations than iterative methods. Fortunately, recent advances in surrogate modeling techniques such as GPEs can negate the computational expense by providing a low-cost substitute for the biophysics model during the fitting process. Instead of performing well over a million biophysics model simulations, one can replace these with computationally inexpensive surrogate models such as GPEs, as used here, which can be conveniently trained on the results of just over a hundred biophysics model evaluations. Even though our biophysics model is already extremely fast (seconds vs. hours) compared to other cardiac mechanics-based growth models ^11–15^, GPEs significantly accelerated model calibration, resulting in a two-step optimization pipeline that took only 10 minutes on a desktop computer. While GPEs do not necessarily possess the same predictive capacity as neural networks ^42^, they require fewer data to calibrate and return not only the predicted outcome but also its uncertainty inherent to the emulator. Surrogate modeling also enabled performing a global sensitivity as part of the calibration pipeline that helped us eliminate parameters that were not relevant to this model optimization. A global sensitivity analysis escalated by GPEs is prevalent in another study of a finite-element cardiac electromechanical model ^43^.

One particular challenge we encountered while developing our method is how to handle parameter sets that result in unstable biophysics model outcomes. For example, the Newton-Raphson method that we employ to solve for the cardiac geometry in our model tends to diverge when there is a mismatch between the total stressed blood volume (*SBV*) and the ventricular wall area (*A*_m,ref_). Initially, we directly used all biophysics model outputs for GPE training, which led to several poorly trained emulators. We addressed this by implementing two filtering steps: first, using multivariate Mahalanobis filtering of the biophysics model outputs to exclude outliers prior to GPE training and, second, exclude parameter sets that resulted in GPE outputs that were non-physiological, e.g. MAP < 25 mmHg and > 250 mmHg. While this could merely be an issue for our biophysics model, we recommend others who want to use this method to implement similar filtering steps. We included these filtering functions in the GitHub repository.

### 4.3 Limitations

Our coupled biophysics and machine learning models to predict cardiac growth probability have several limitations. The first limitation is our biophysics model’s simplified representation of certain cardiovascular physiology. These simplifications were chosen to reduce computational power and time, enabling it to simulate 3 months of cardiac growth in about one second. The main simplification is in the cardiac geometry, which we modeled using the TriSeg method, i.e. simulating the atrioventricular complex using thick-walled spherical segments instead of a full 3D finite-element approach. Moreover, the vasculature is simulated using a 0-D electrical analog model, instead of a more comprehensive fluid mechanics approach. While we and others have demonstrated that these simplifications do not impede such a model’s ability to predict cardiac physiology ^28^ and growth ^16,17^ following volume overload, and previously pressure overload, cardiac dyssynchrony, and myocardial ischemia it could potentially prohibit accurate cardiac growth predictions for other diseases and treatments where spatial heterogeneity in the cardiac wall are important.

Another biophysics model limitation is that hemodynamic reflexes such as the baroreceptors were not considered. This could significantly affect the accuracy of our results and could have been the cause of the underestimate of predicted LVEDWth (Fig. 10). It was recently shown that including baroreflexes and/or sympathetic vs. parasympathetic regulation increases the accuracy of ventricular growth predictions ^44,45^. In our proposed two-step framework, calibration of parameters for such reflex models would need to be included in both steps depending on the time scale: baroreflexes act on the order of seconds so will need to be included in the first step, whereas mechanisms such as the renal system act in hours/days and will need to be included in the second calibration step.

A limitation of the GPEs was the lack of output interaction in the surrogate models since we used a separate GPE for each model output. For example, end-diastolic and end-systolic pressure and volume are related to one another through the passive and active myocardial stiffness (Eqs. 3, 4). We attempted to train a multi-variate GPE to account for these interactions, however, we failed to accurately train this emulator to our biophysics model and instead employed single-variate output GPEs as in Coveney et al. ^24^ and Longobardi et al. ^25^. In the future, we will utilize dimension reduction strategies (e.g. principal component analysis) to obtain independent outputs for GPE training.

Finally, it is important to consider the initial input parameter ranges. When too small the parameter space does not include the optimal region; when too large, the number of unstable simulations can be detrimental to the GPE training process and cause large model uncertainty. However, this issue is even more persistent in iterative methods, where one typically has to choose a single initial parameter estimate; an initial choice positioned too far away from the global minimum could lead to finding a local instead of global minimum of the objective function. At least in BHM, one has to choose a range of initial parameter choices, which substantially reduces the risk of not including the optimal parameters.

### 4.4 Future perspectives

With the increased availability of comprehensive diagnostic data, cardiac models are inching closer to realizing a precision medicine approach. It is vitally important for both clinicians and researchers to be aware of the potential error within the results of patient-specific models to make the most informed decisions. The combined biophysics and machine learning modeling framework we proposed in this study can be easily translated to a routine clinical environment. It can be optimized to data from a single patient in under 10 minutes on a standard computer, and is capable of predicting long-term cardiac growth probability following different therapeutic interventions in several seconds once calibrated.

While in this investigation we tested and validated our framework to means and standard deviations of previously published canine data (Fig. 10), these methods can be readily used to calibrate our model to data from an individual patient. In lieu of the standard deviation of a cohort, one can use inter-operator variability in MRI analysis, such as proposed by Rodero et al. ^26^, to identify all possible growth outcomes for a single patient. This data uncertainty will be propagated into the long-term growth predictions. For instance, using data from low-resolution ultrasound will likely lead to a larger possible range of growth outcomes than data from high-resolution CT images. We demonstrated this in our synthetic cases using 10*%* and 20*%* standard deviations, where with more uncertainty in the target data, a wider range of growth outcomes is simulated (Fig. 8).

In conclusion, our study marks a significant stride in cardiac growth model calibration and predictions. Instead of providing a single growth prediction that will almost certainly be incorrect, these methods could provide a range of potential patient-specific, long-term outcomes to clinicians.

## 5 Data availability

These methodologies and results will be made publicly available on GitHub upon publication of this paper at http://github.com/BeatLabUCI/. Curated Jupyter notebooks with extensive documentation are included to reproduce all results in this manuscript, as well as tutorial notebooks to help others use our code to calibrate their own models. Further inquires can be addressed to the corresponding authors.

## Acknowledgments

This research was funded by the National Institutes of Health (R01HL159945).

## Authors contributions

Author 1: Methodology, Investigation, Formal analysis. Author 2: Methodology, Investigation, Formal analysis, Funding acquisition, Supervision.

## Appendix A

Emulators were trained on 80% of the simulation data, and independently validated to the remaining 20%. Emulator quality was similar across all cases, Fig. A1 is showing the validation of the emulators trained during the final wave of the *σ*_20%_ synthetic baseline calibration case.

**Figure A1:**
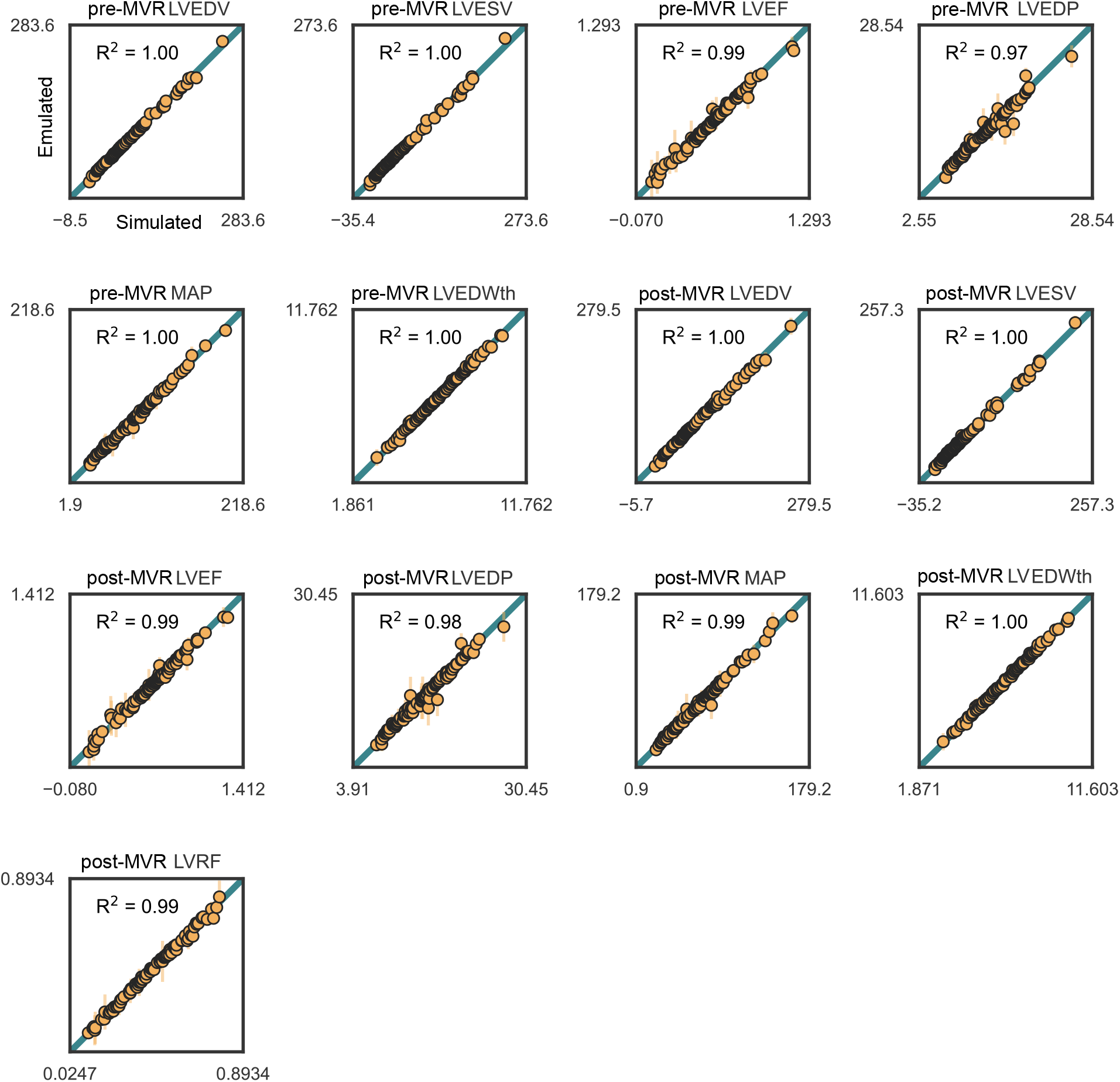
GPE validation comparing emulated versus simulated outputs. To validate GPE accuracy, simulated output values (horizontal axis) were compared to emulated values (vertical axis). Emulators were trained on 80% of the simulation data, and independently validated to the remaining 20%. The emulator quality was similar across all case, with the validation shown here belonging to the baseline phase of the *σ*_20%_ synthetic case,

## Appendix B

Analysis of the two-step approach methodology. Fig. B1 shows that randomly recombining the calibrated baseline and growth parameters for the baseline phase of the *σ*_10%_ synthetic case does not have a substantial impact on the final growth simulations, with only LVEDV slightly underpredicted. Fig. B2 shows that concurrently fitting the baseline and growth parameters results in growth simulations that do not cover the full range of possible outcomes, in contrast to our two-step method. These results combined indicate the usefulness of our two-step approach.

**Figure B1:**
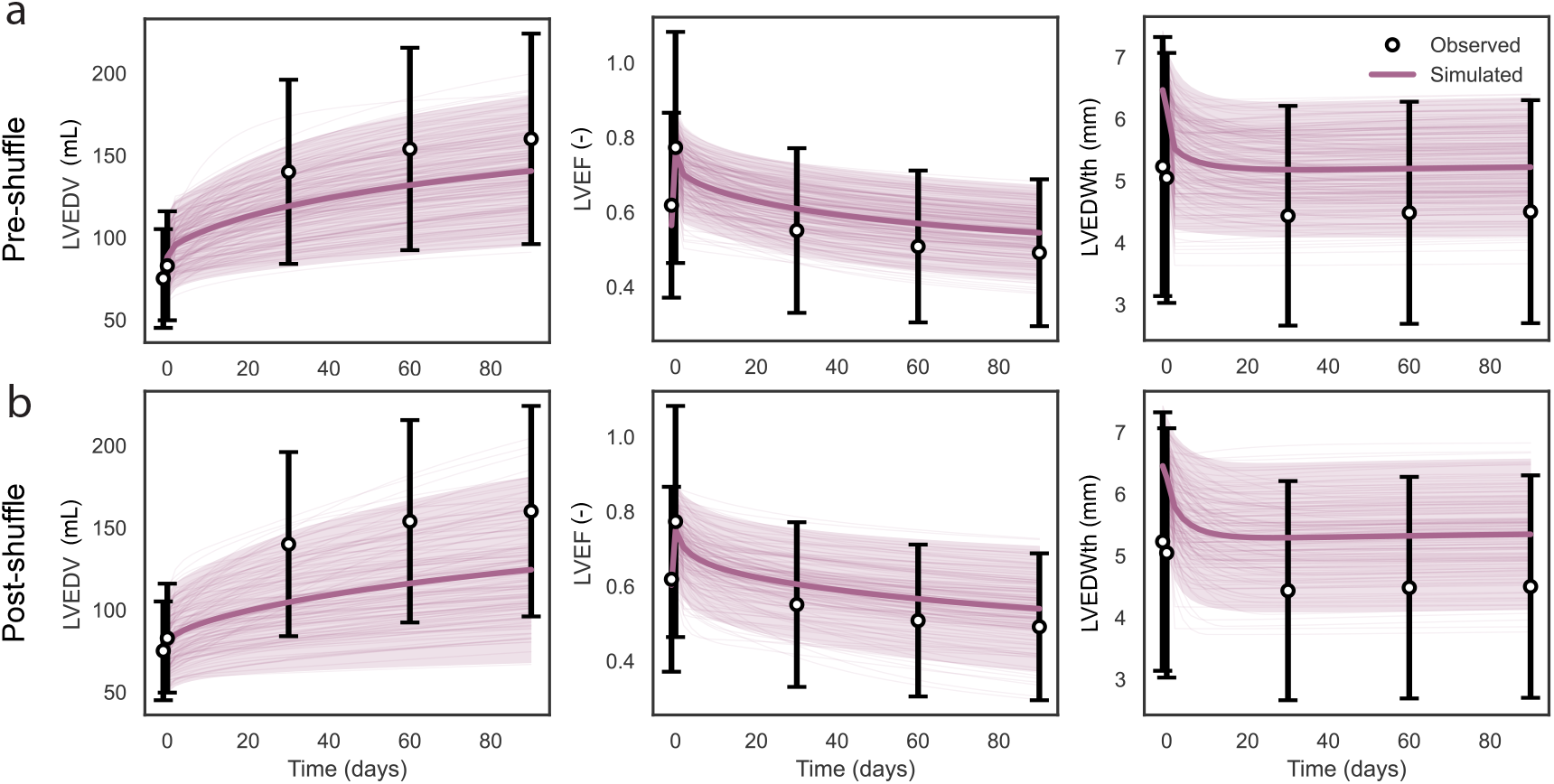
Recombination of growth parameters sets and baseline fits. Shuffling growth parameter sets with baseline fits is possible with this study’s independent two-step BHM calibration method. The predictions do not drastically change between the pre-shuffle (a) and post-shuffle (b) growth simulations.

**Figure B2:**
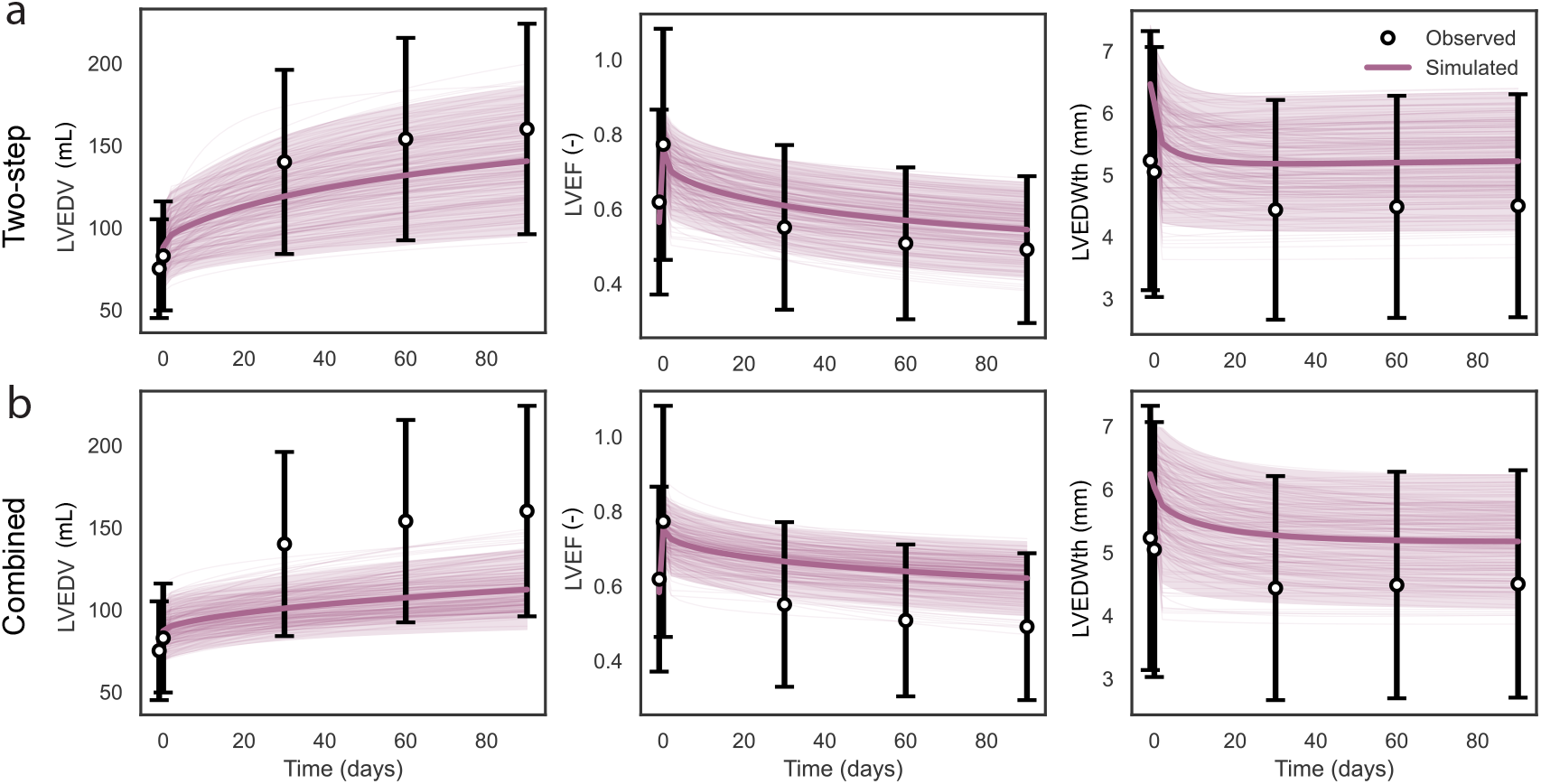
Two-step fitting versus combined baseline and growth fitting. Our two-step calibration approach (here shown for the synthetic *σ*_20%_ case) resulted in growth simulations with a variance comparable to the data (a), whereas concurrently calibrating baseline and growth parameters resulted in growth simulations with a much lower variance.

